# Identification of mitochondria and macrophage activation related hub genes in sarcopenia

**DOI:** 10.1101/2025.06.04.657824

**Authors:** Yanjin Wang, Shan Gu, Yang Li, Yingjie Zhou

## Abstract

**BACKGROUND:** To explore hub genes and pathways related to sarcopenia progression associated with mitochondria and macrophage activation.

**METHODS:** GEO datasets GSE8479, GSE1428, and GSE136344 were obtained based on GEO database. MRGs and MARGs were acquired. Thereafter, GO, KEGG, GSEA and PPI analysis were implemented. Later, pooled GEO datasets (Combined Datasets) were further obtained to construct the sarcopenia diagnostic model and sarcopenia subtypes.

**RESULTS:** A total of 62 M&MARDEGs were finally identified. There were altogether 9 model genes incorporated into the multifactorial regression model. Immunoinfiltration analysis revealed low scores of M&MARDEGs in the samples with sarcopenia. For macrophage activation in sarcopenia samples, most immune cells were strongly correlated in LowScore group (r = -0.707, p < 0.05). Many immune cells in Mitochondria & Macrophage Activation (HighScore) group were strongly correlated with each other (r = -0.661, p < 0.05).

**CONCLUSION:** Screening key genes for sarcopenia associated with mitochondrial and macrophage activation in this study assists in understanding the pathogenesis of sarcopenia and providing new targets for intervention.

## 1. Introduction

Sarcopenia, as the skeletal muscle aging disorder, constitutes a significant health burden for the patient and the community [1–3]. Its prevalence among people aged 60 and above is 5.7% to 23.9%. Skeletal muscle mass loss, muscle strength decline, and behavioral deficits are the three main characteristics of sarcopenia, which directly impairs patients’ quality of life. The current management of sarcopenia consists of exercise intervention and nutritional supplementation [4]. However, effective molecular markers for early diagnosis and personalized treatment are lacking.

Mitochondria are the energy factories in the cell and their functional state directly affects metabolic activities and physiological functions in the cell. Mitochondrial dysfunction is recognized as one of the key mechanisms in the pathogenesis of sarcopenia [5]. Skeletal muscle cells have extremely high energy requirements and are highly sensitive to mitochondrial dysfunction. Mitochondrial dysfunction directly impairs the physiological function of skeletal muscle [6–7]. Also, mitochondrial dysfunction is recognized to be an important hallmark for cellular senescence [8]. Accumulation of mitochondrial DNA (mtDNA) damage during aging weakens mitochondrial self-repair and aggravates mitochondrial dysfunction. In addition, mitochondrial dysfunction disrupts intracellular calcium homeostasis and muscle protein metabolic homeostasis.

Research data have shown that macrophage is known to regulate inflammation by promoting aggregation of inflammatory factors [9–10]. Macrophage activation mainly consists of transformation between M1 macrophages and M2 macrophages. When skeletal muscle atrophy occurs, macrophages transform from M2 to M1 by increasing the inflammatory factor levels like interleukin (IL)-6 or tumor necrosis factor (TNF)-α [11–12]. Therefore, macrophage activation in skeletal muscle is important for muscle atrophy and helps improve skeletal muscle atrophy through modulating the M1-M2 macrophage balance.

To further elucidate the role of mitochondrial function and macrophage activation in the process of sarcopenia and the related molecular mechanisms, various bioinformatics analysis approaches, including batch effect correction, differential expression analysis, enrichment analysis, and machine learning model construction, were utilized in the present work, aiming to reveal the molecular features associated with sarcopenia and establish the diagnostic model for improving clinical application.

## 2. Materials and Methods

### 2.1 Data acquisition

The R package GEOquery[13] (Version 2.70) from GEO database[14] (https://www.ncbi.nlm.nih.gov/geo/) was used to download muscle disease (sarcopenia) datasets GSE8479[15], GSE1428[16] and GSE136344[17]. The samples of the datasets GSE8479, GSE1428 and GSE136344 were all derived from homo sapiens and the tissue sources were all lateral thigh muscle tissues. For details, see Table 1.

The GSE8479 dataset had the chip platform of GPL2700, and it consisted of 25 sarcopenia samples as well as 26 control samples. For the GSE1428 dataset, its chip platform was GPL96, containing 12 sarcopenia samples together with 10 control samples. Dataset GSE136344 had a chip platform of GPL5175 and contained 12 sarcopenia samples as well as 11 control samples. The present work enrolled both sarcopenia and control groups.

Mitochondrial genes can be acquired in GeneCards database[18] (https://www.genecards.org/), which offers integrated data regarding human genes. Mitochondria were utilized to be the search keyword, with MRGs satisfying "Protein Coding" and "Relevance Score > 1" (n = 3,503) being collected. Additionally, "mitochondria" were made as a keyword during search at PubMed website (https://pubmed.ncbi.nlm.nih.gov/) to get MRGs (n =25) at publications[19]. Following combining and de-repeating, there were altogether 3509 MRGs acquired. for details, see Table S1.

We searched by the term "macrophage activation" through GeneCards database to collect MARGs. In total, 417 MARGs were obtained by retaining only those satisfying "Protein Coding" and "Relevance Score > 1". Moreover, "Macrophage Activation" was made as a keyword for search at PubMed website (https://pubmed.ncbi.nlm.nih.gov/) to get MARGs at publications[20], and five MARGs were acquired. There were 418 MARGs discovered following the combination and duplicate removal. For more details, see Table S2. Finally, we obtained 188 sarcopenia-related Mitochondria & Macrophage Activation-Related Differentially Expressed Genes (M&MARDEGs) by overlapping 3509 MRGs and 418 MARGs. For the detailed information, see Table S3.

R package sva [21] (Version 3.50.0) is adopted for de-batching GSE8479 and GSE1428 to get an integrated GEO data set (Combined Datasets), which covered 37 sarcopenia and 36 control cases. At last, R package limma [22] (Version 3.58.1) was utilized for normalizing Combined Datasets, followed by normalization of annotation probe. Principal component analysis (PCA) was performed for verifying the expression matrix before and following batch effect removal[23]. PCA can be applied in reducing data dimensionality, and it is capable of extracting feature vector (component) data among high-dimensional data and convert them in low-dimensional counterparts and uses 2D or 3D graphs to show these features.

### 2.2 Sarcopenia related mitochondria & macrophages activate related differentially expressed genes

All samples were classified as a sarcopenia or a control group, respectively, according to sample grouping in the integrated GEO data set. Differences in genes in the sarcopenia and control groups were analyzed using R-packet limma (Version 3.58.1). The thresholds for |logFC| > 0 and p value < 0.05 were set as differentially expressed genes (DEGs). Genes with logFC > 0 and p value < 0.05 were up-regulated genes. And Genes with logFC < 0 and p value < 0.05 were down-regulated genes. The Benjamini-Hochberg (BH) approach was used to correct p-value. R package ggplot2 (Version 3.4.4) was applied in mapping our differential analysis results.

In order to obtain M&MARDEGs, the intersection of DEGs in integrated GEO data set with M&MARGs was taken, followed by the drawing of Venn diagram. M&MARDEGs were obtained, and the Top20 M&MARDEGs were mapped using R-package heatmap (Version 1.0.12).

### 2.3 Gene Ontology (GO) Functional enrichment analysis, Kyoto Encyclopedia of Genes and Genomes (KEGG) enrichment

GO enrichment[24] has been commonly used to conduct large-scale functional annotation studies, consisting of Biological Process (BP), Cell Component (CC), and Molecular Function (MF). The KEGG database[25] has been commonly used in storing data regarding biological pathways, genomes, drugs, and diseases, among other things. R-pack clusterProfiler[26] (Version 4.10.0) was employed for GO and KEGG analyses on M&MARDEGs. The selection criteria included p < 0.05 and FDR (q) < 0.05, and BH approach was adopted to correct p-value.

### 2.4 Gene Set Enrichment Analysis (GSEA)

GSEA[27] can be used for assessing gene distribution tendency with pre-defined gene sets among genes ranked by phenotypic correlation, thus judging the gene contributions to phenotype. The present work sequenced genes from a combined GEO dataset based on logFC values in the sarcopenia versus control groups. Thereafter, R-package clusterProfiler (Version 4.10.0) was utilized for analyzing each gene from the integrated GEO data set. All parameters used in GSEA included: Seeds for 2020, and every gene set contains 10-500 500 genes. Meanwhile, c2 gene sets were obtained by using Molecular Signatures Database (MSigDB). All. V2023.2. Hs. Symbols for GSEA, Our GSEA criteria were p < 0.05 and FDR (q-value) < 0.05, and p-value was corrected by BH approach.

### 2.5 Establishment of the sarcopenia diagnostic model

For obtaining the Sarcopenia diagnostic model from those combined GEO Datasets, M&MARDEGs were subjected to logistic regression. When dependent variables were binary, i.e. sarcopenia and control groups, the relation of independent variable with dependent variable was examined through logistic regression. With p value < 0.05 as the criterion, M&MARDEGs were screened, and a logistic regression model was constructed.

Then, LASSO (Least Absolute) was performed by R package glmnet[28] using set.seed (500) and family= "binomial" as parameters based on M&MARDEGs included in the logistic regression model Shrinkage and Selection Operator regression analysis. Based on linear regression, LASSO regression analysis could reduce model overfitting while improving model generalizability through introducing one penalty term (|lambda x slope|). Both diagnostic model and variable locus diagrams can be drawn for visualizing LASSO regression. In this study, our LASSO regression results were a sarcopenia diagnostic model with M&MARDEGs associated with mitochondrial & macrophage activation as Model genes. At last, according to risk coefficients in LASSO regression, LASSO risk Score (RiskScore) values were determined below:

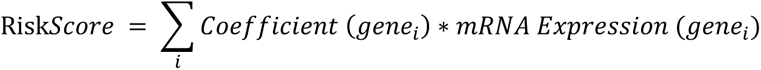

Thereafter, on the basis of M&MARDEGs included into our logistic regression model, the an SVM (Support Vector Machine) algorithm was employed for constructing a SVM model [29]. M&MARDEGs were chosen in line with gene number showing greatest accuracy whereas smallest error rate. We selected the genes obtained from the intersection of M&MARDEGs that were screened by LASSO and SVM analyses in later analyses.

### 2.6 Verification of our as-constructed diagnostic model for sarcopenia

Then, rms was applied in constructing the Nomogram for M&MARDEGs. A Nomogram is comprised by various disjoint line segments representing functional relations of independent variables in the planar rectangular coordinate system. According to multifactor regression analysis, we set certain scales to characterize the risk of all variables incorporated into the multifactor regression model. Eventually, total risk was calculated for predicting the even probability.

Calibration graph is plotted for evaluating the model performance in predicting real result through drawing model-predicted and real probabilities in diverse situations. It can be utilized to analyze model fitting using Logistic regression and real situation. We use variables in our multifactor regression model to be key genes in later analyses.

Decision curve analysis (DCA) can be utilized for evaluating clinical predictive models, molecular markers and diagnostic tests. Finally, R package ggDCA was employed for drawing DCA plots for evaluating the Logistic regression model resolution and accuracy.

In addition, R package pROC was utilized for plotting ROC Curve in an integrated GEO data set and calculating the Area Under the ROC Curve (AUC). To evaluate Linear Predictors (Logistic regression models) for their diagnostic effect on sarcopenia development, the AUC is usually 0.5-1, with a value approaching 1 indicating the superior diagnosis.

### 2.7 Protein-Protein Interaction (PPI) Network

Protein-protein interaction networks are composed of individual proteins through their interactions with each other. The STRING database[30] is a database that searches for interactions of known proteins with predicted ones. We used the STRING database, constructed PPI networks through setting human as the biological species and confidence ≥ 0.150 pairs of key genes, and employed Cytoscape for visualizing the PPI network model.

The GeneMANIA website[31] was utilized for predicting functionally-similar key genes and constructing the interaction network.

### 2.8 Construction of regulatory network

miRNA has a key effect on regulating biological occurrence and progression. It can modulate several target genes, meanwhile, one target gene is modulated via several miRNAs. For analyzing the relation of M&MARDEGs with miRNAs, TarBase[32] database (http://www.microrna.gr/tarbase) was used for searching M&MARDEGs related miRNAs. In addition, we visualized the mRNA-miRNA network using Cytoscape. Transcription factor (TF) can modulate gene levels in the post-transcriptional phase through interaction with M&MARDEGs. Through utilizing ChIPBase database[33] (http://rna.sysu.edu.cn/chipbase/) for retrieving TF, TF analysis on M&MARDEGs was conducted, and later, Cytoscape[34] was applied in visualizing the mRNA-TF network.

### 2.9 Analysis of key DEGs

For identifying the possible mechanism, relevant biological features and pathways underlying DEGs related to sarcopenia, we conducted Mann-Whitney U test (Wilcoxon rank sum test) according to key genes in different sarcopenia (subgroup: sarcopenia) and normal (subgroup: control) data sets, and R package ggplot2 was utilized for drawing comparison plots to display differential analysis results.

Then, key genes were selected according to differential expression results, while ROC curves for screened key genes from different data sets were examined, and the results were displayed. As the coordinate schema analysis approach, ROC Curve [35] is utilized for choosing the optimal model, discarding the suboptimal model, or setting the best threshold in a model. The ROC curve can comprehensively reflect whether continuous variables were sensitive and specific. Typically, the composition approach can reflect the association of sensitivity with specificity. The AUC value is usually 0.5-1, with a value approaching 1 suggesting superior diagnostic performance. The AUCs of 0.5-0.7, 0.7-0.9 and >0.9 show low, certain and high accuracy, separately. The R package proc was applied in plotting ROC curves for screened key genes from diverse datasets, and AUC values were determined for assessing the diagnostic performance of expression of key genes in survival in patients with sarcopenia.

### 2.10 Construction of high-low rating grouping of mitochondrial & macrophage activation

Single-Sample gene-set Enrichment Analysis (ssGSEA) [36] quantifies the gene relative level in the dataset sample. We utilized ssGSEA algorithm for calculating Mitochondria & Macrophage Activation Score (M&MA.Score) in each sample with R package GSVA, according to the key gene expression matrix in the Combined Datasets. In addition, median expression of mitochondrial & macrophage activation score (M&MA.Score) divided the sarcopenia sample in the high (HighScore) or low-score (LowScore) group.

### 2.11 Immunoinfiltration Analysis (ssGSEA)

All infiltrated immune cell types, e.g. Activated CD8+ T cell, Activated dendritic cell, Gamma-delta T cell, Natural killer T cell, and Regulatory T cell, were labeled. Subsequently, ssGSEA was adopted for determining enrichment scores, which represented the immune cell infiltration level of every sample, and subsequently the immune cell infiltration matrix of Combined GEO datasets was obtained. Later, group comparison plots were drawn with R-package ggplot2 (Version 3.4.4) for presenting differential immune cell levels in different groups of disease groups in Combined GEO Datasets. Subsequently, significantly differential immune cells in two groups were selected in later analyses, the associations of immune cells were determined using Spearman algorithm, with the correlation heatmap being plotted with R pheatmap for illustrating immune cell correlation analysis results. Correlations of key genes with immune cells were explored with Spearman algorithm, whereas the correlation bubble map was prepared with R package ggplot2 (Version 3.4.4) for displaying the key gene-immune cell correlation.

### 2.12 Construction of subtypes of sarcopenia

As the resampling-based algorithm, Consensus Clustering[37] identifies every member as well as the subgroup number and later verifies the clustering rationality. It is the multiple iterations of a subsample dataset that induces sampling variability using subsampling to provide indicators for parametric decision making and cluster stability. It uses R package ConsensusClusterPlus[38] (Version 1.62.0) to identify the different disease subtypes of a sarcopenia sample in the Combined Datasets on the basis of key genes. The cluster number is set at 2-9, 80% of total sample is drawn 50 times, clusterAlg = "km", distance = "pearson". Then, the expression difference in key genes among diverse sarcopenia disease subtypes from the integrated GEO data set was analyzed by expression value heat maps. And further verified differences in key gene expression among diverse disease subtypes by subgroup comparison maps.

### 2.13 Statistical analysis

R software (Version 4.2.2) was employed to process and analyze data. In addition, the independent Student T Test was used to compare normally-distributed continuous variables in two groups. And non-normally-distributed variables were analyzed by Mann-Whitney U Test, namely Wilcoxon Rank Sum Test. Among-group differences were analyzed by Kruskal-Wallis test. Correlation coefficients among diverse molecules were examined through Spearman correlation analysis. If not specified, p-value was bilateral, and p < 0.05 stood for significant differences.

## 3. RESULTS

**Technology roadmap:**

**Fig 1.**
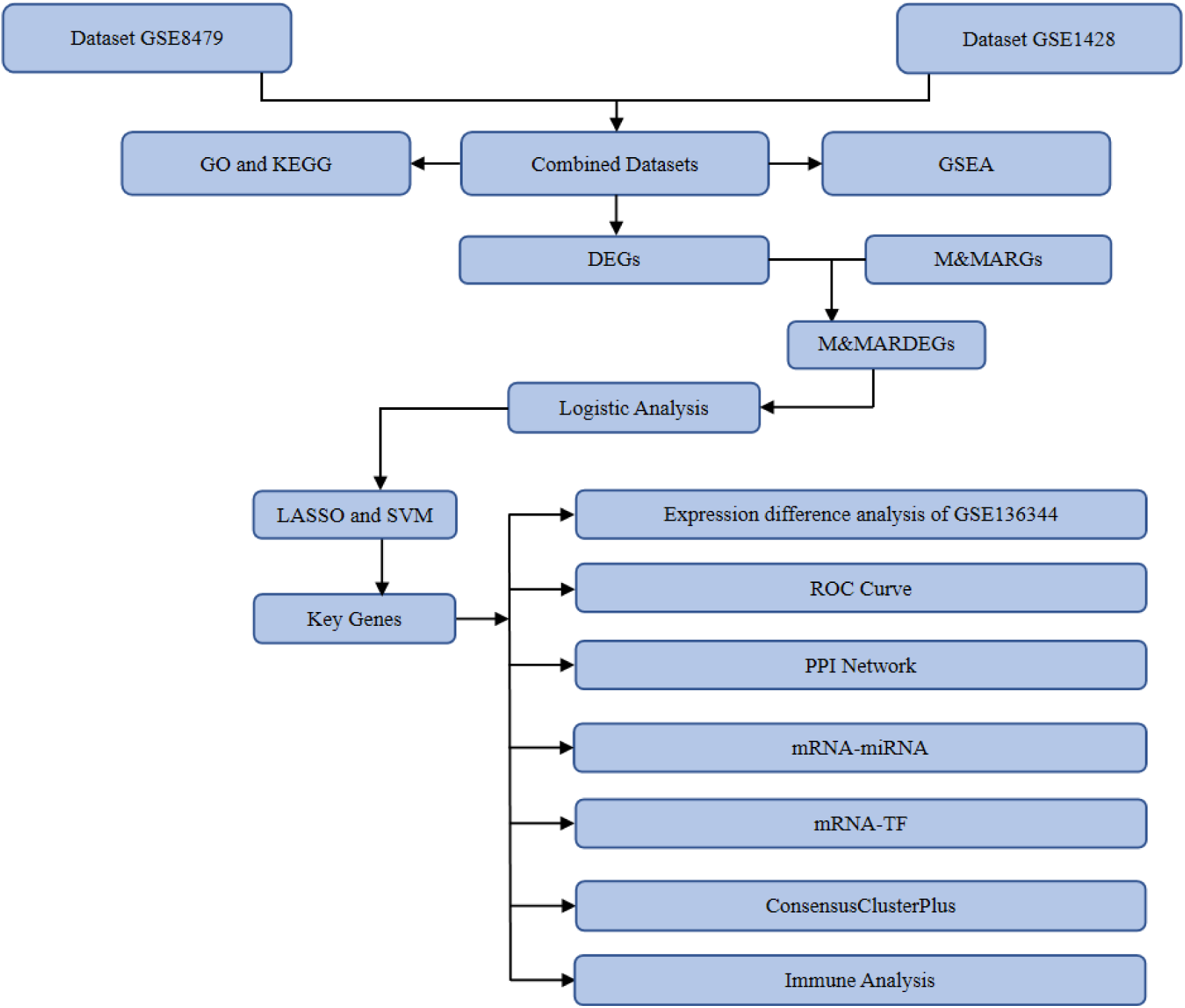
Technology Roadmap. GO: Gene Ontology. KEGG: Kyoto Encyclopedia of Genes and Genomes. GSEA: Gene Set Enrichment Analysis. DEGs: Differentially Expressed Genes. M&MARGs: Mitochondria & Macrophage Activation - Related Genes. M&MARDEGs: Mitochondria & Macrophage Activation - Related Differentially Expressed Genes. LASSO: Least Absolute Shrinkage and Selection Operator. SVM: Support Vector Machine. ROC: Receiver Operating Characteristic. PPI: Protein-Protein Interaction. TF: Transcription Factor.

### 3.1 Data download and calibration

First, both GSE8479 and GSE1428 Sarcopenia Datasets were treated with R-packet sva for removing batch effect. Later, the integrated GEO data set was obtained. Subsequently, the differences in data set expression before and following removing batch effect were analyzed using Fig 2A-B distribution box plots. Secondly, the low-dimensional feature distribution of data sets before and following batch effect removal was examined through PCA (Fig 2C-D). As revealed by distribution box and PCA plot analyses, batch effects for samples from the Sarcopenia dataset were nearly totally removed.

**Fig 2.**
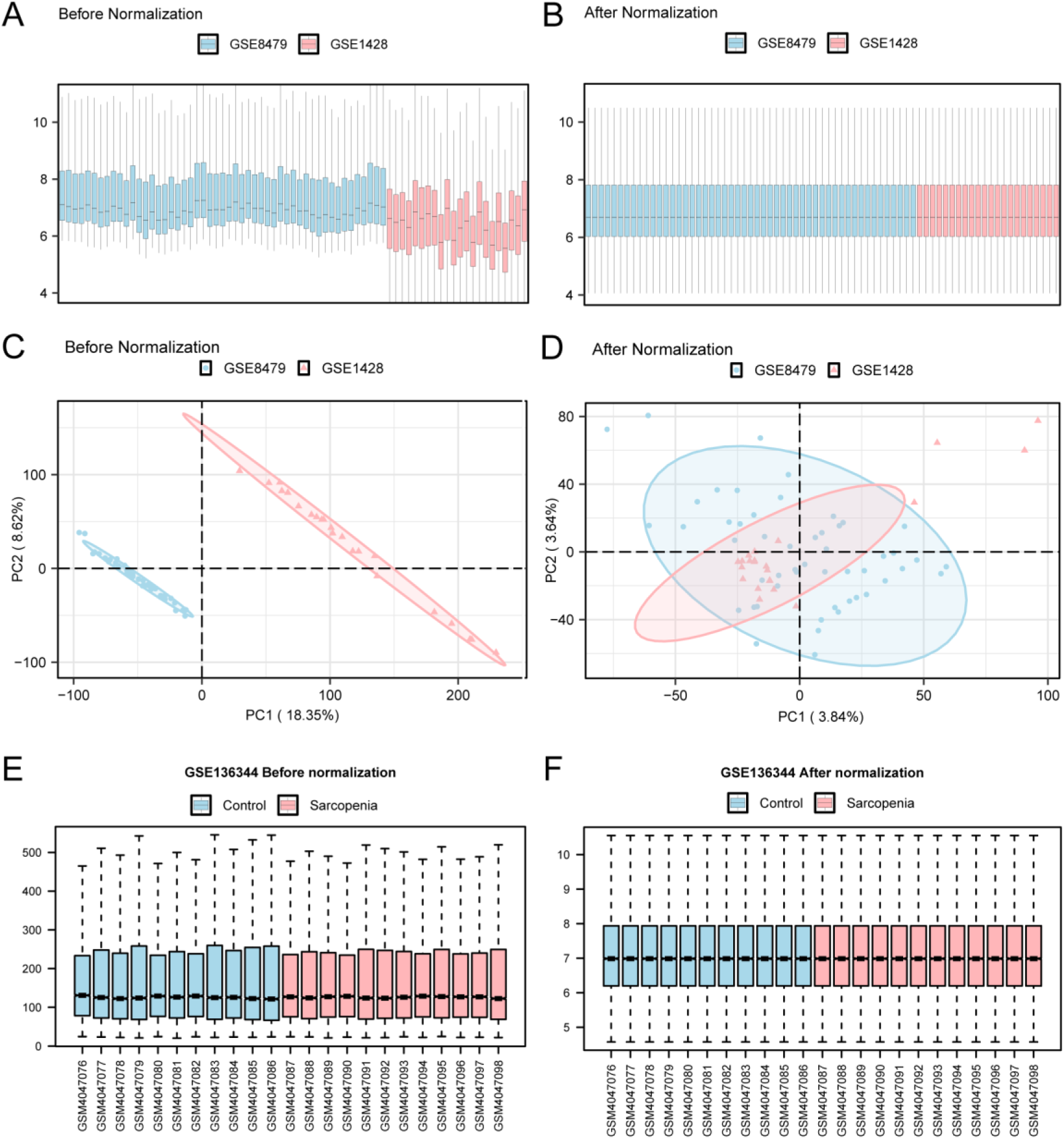
Removal of Dataset Batch Effects. Distribution boxplots for Combined Datasets of GEO before (A) and following (B) removing batch. PCA plots for Combined Datasets of GEO before (C) and following (D) removing batch. Box diagrams showing gene level distribution in GSE136344 samples before (E) and after (F) correction. PCA, Principal Component Analysis; Blue for sarcopenia dataset GSE8479, pink for sarcopenia dataset GSE1428, pink for sarcopenia dataset GSE136344, blue for control dataset.

Samples from GSE136344 dataset were classified as sarcopenia group or control group. After normalizing the GSE136344 dataset (Fig 2E-F), the data cleaning including the probe was annotated, with data distribution before and following normalization being plotted with a box plot. After normalization, diverse samples almost had the same expression levels in the dataset.

### 3.2 Mitochondrial & macrophage activation-related differentially expressed genes associated with sarcopenia

Samples in Combined GEO Datasets were classified as a sarcopenia group or a control group. For analyzing differential gene levels in sarcopenia versus control groups of integrated GEO dataset (combined Datasets), difference analysis of the combined GEO data set was conducted using limma package R to obtain DEGs. There were altogether 3384 DEGs satisfying |logFC| > 0 and p < 0.05 in the integrated GEO dataset, including 1707 with up-regulation (logFC > 0 and p < 0.05) and 1677 with down-regulation (logFC < 0 and p < 0.05). From our difference analysis of this dataset, the volcano map was drawn (Fig 3A).

**Fig 3.**
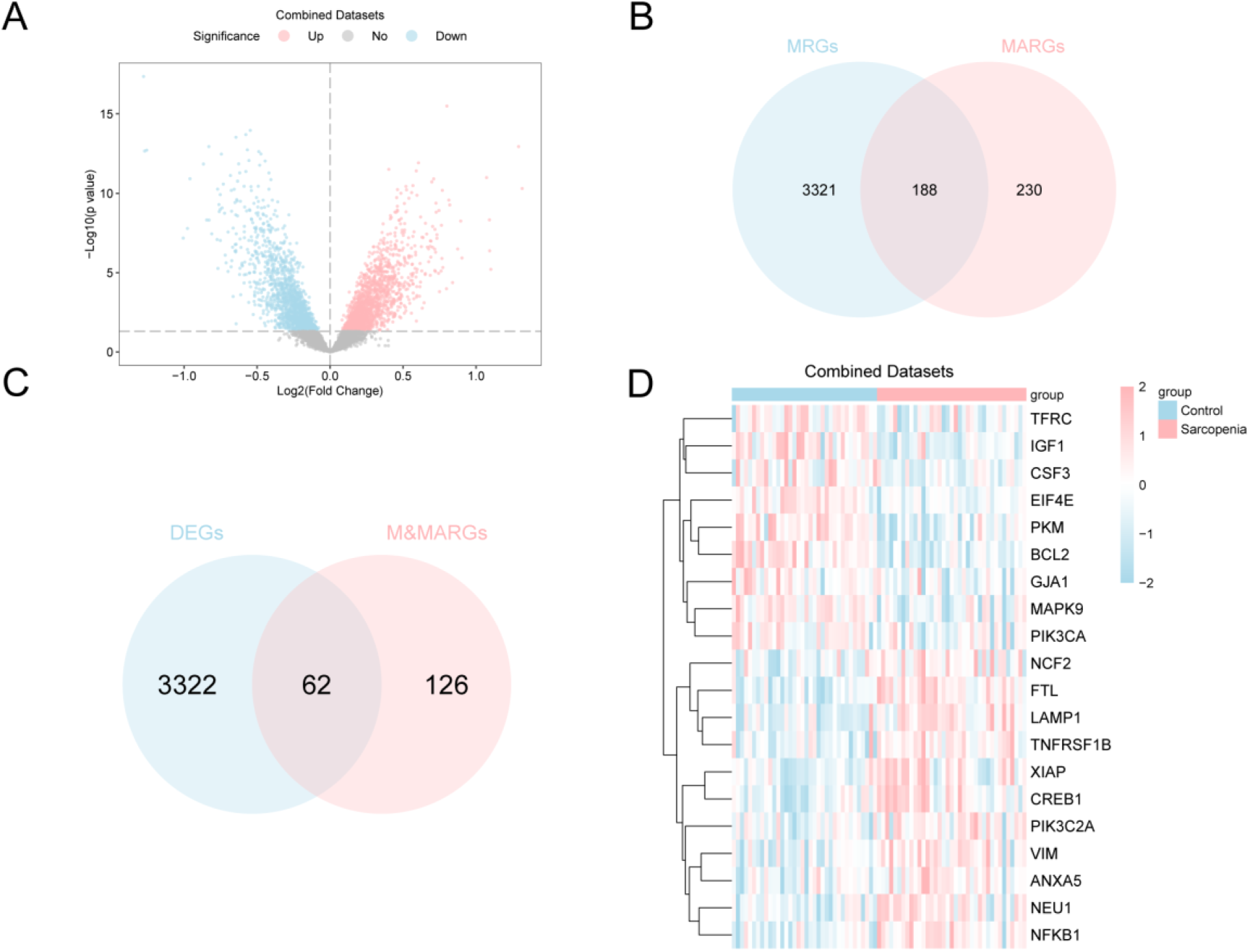
Differential Gene Expression Analysis. A. Differential gene expression analysis volcano map in the Sarcopenia versus Control groups in Combined GEO Datasets. B. Venn map showing mitochondria-associated genes (MRGs) and macrophage Activation associated genes (MARGs). C. Venn diagram showing DEGs and M&MARGs in the Combined Datasets. D. Heat map showing mitochondria & Macrophage activation-related differentially expressed genes (M&MARDEGs) in the Combined Datasets. MRGs: Mitochondria-Related Genes. MARGs: Macrophage Activation-Related Genes. DEGs: Differentially Expressed Genes; M&MARGs: Mitochondria & Macrophage Activation - Related Genes. M&MARDEGs: Mitochondria & Macrophage Activation - Related Differentially Expressed Genes. Sarcopenia (pink) and Control (blue). Pink and blue colors suggest high and low expression separately.

For obtaining M&MARDEGs, we first intersected 3509 mitochondria-related genes (MRGs) and 418 macrophage activation-related genes (MARGs) and drew Venn map (Fig 3B). A total of 188 Mitochondria & Macrophage Activation-Related Genes (M&MARGs) were obtained. Then, all the obtained |logFC| > 0 and p < 0.05 DEGs and mitochondria & Macrophage activation-related genes (M&MARGs) were mitochondria and macrophage activation related genes (Fig 3C). A total of 62 M&MARDEGs related to mitochondrial & macrophage activation were obtained, which were *FTL, BCL2, PKM, LAMP1, NFKB1, IGF1, VIM, NEU1, MAPK9, CREB1, TNFRSF1B, EIF4E, XIAP, HSPD1, FTL, BCL2, PKM, LAMP1, NFKB1, IGF1, VIM, CREB1, TNFRSF1B, Eif4e, XIAP, HSPD1, respectively. IRAK1, SPHK2, BMP6, MTOR, TF, NCF2, PPARA, PARP1, HSPA4, MAPK3, TUG1, PIK3C2A, ANXA5, VEGFA, CSF3, STX11, MAPK8, CD4, SRC, TFRC, NOS2, RELA, TXNRD1, IKBKG, JUN, GJA1, SLC7A7, PLA2G4A, STAT3, IRF1, AKT2, NOTCH1, MAPK10, PIK3R1, CXCL8, PTK2B, MCM4, FAS, TNFRSF1A, AP3B1, PIK3CA, NAGA, CD36, CASP3, PRKCE, MPO, ICAM1, PRKCD.* According to the intersection results, samples in the integrated GEO dataset (Combined expression differences in mitochondria & Macrophage activation-related differentially expressed genes (M&MARDEGs) were examined. Later, the heatmaps were drawn by R-packet pheatmap for those top 20 M&MARDEGs (Fig 3D Fig3d).

### 3.3 GO and KEGG analysis

By conducting GO and KEGG analysis, BPs and CCs of 62 mitochondria & macrophage activation-related differentially expressed genes (M&MARDEGs) were further explored. Besides, MF and the relation between KEGG pathways and sarcopenia were analyzed. These 62 mitochondria & macrophage activation-related differentially expressed genes (M&MARDEGs) were conducted GO as well as KEGG analysis (Table 2). Clearly, 62 M&MARDEGs in sarcopenia were mostly related to response to peptide, response to lipopolysaccharide, accharide, response to molecule of bacterial origin, cellular response to biotic stimulus, response to molecule of bacterial origin, cellular response to biotic stimulus, cellular response to lipopolysaccharide and other biological processes (BP); endocytic vesicle, membrane microdomain, membrane raft, cytoplasmic vesicle lumen, secretory granule lumen, and other cell components (CC); MFs including Protein serine/threonine/tyrosine kinase activity, protein serine/threonine kinase activity, the MAP kinase activity, protein serine kinase activity, and phosphatase binding; and KEGG pathways such as Lipid and atherosclerosis, TNF signaling pathway, AGE-RAGE pathway in diabetic complications, Hepatitis B, and Kaposi sarcoma-associated herpesvirus infection. A histogram was plotted to display GO and KEGG results (Fig 4A). Meanwhile, network maps for BPs, CCs, MFs and KEGGs were plotted according to GO and KEGG findings (Fig 4B-E), with lines illustrating annotations of the relevant molecules and items. A larger node indicated more molecules in the entry.

**Fig 4.**
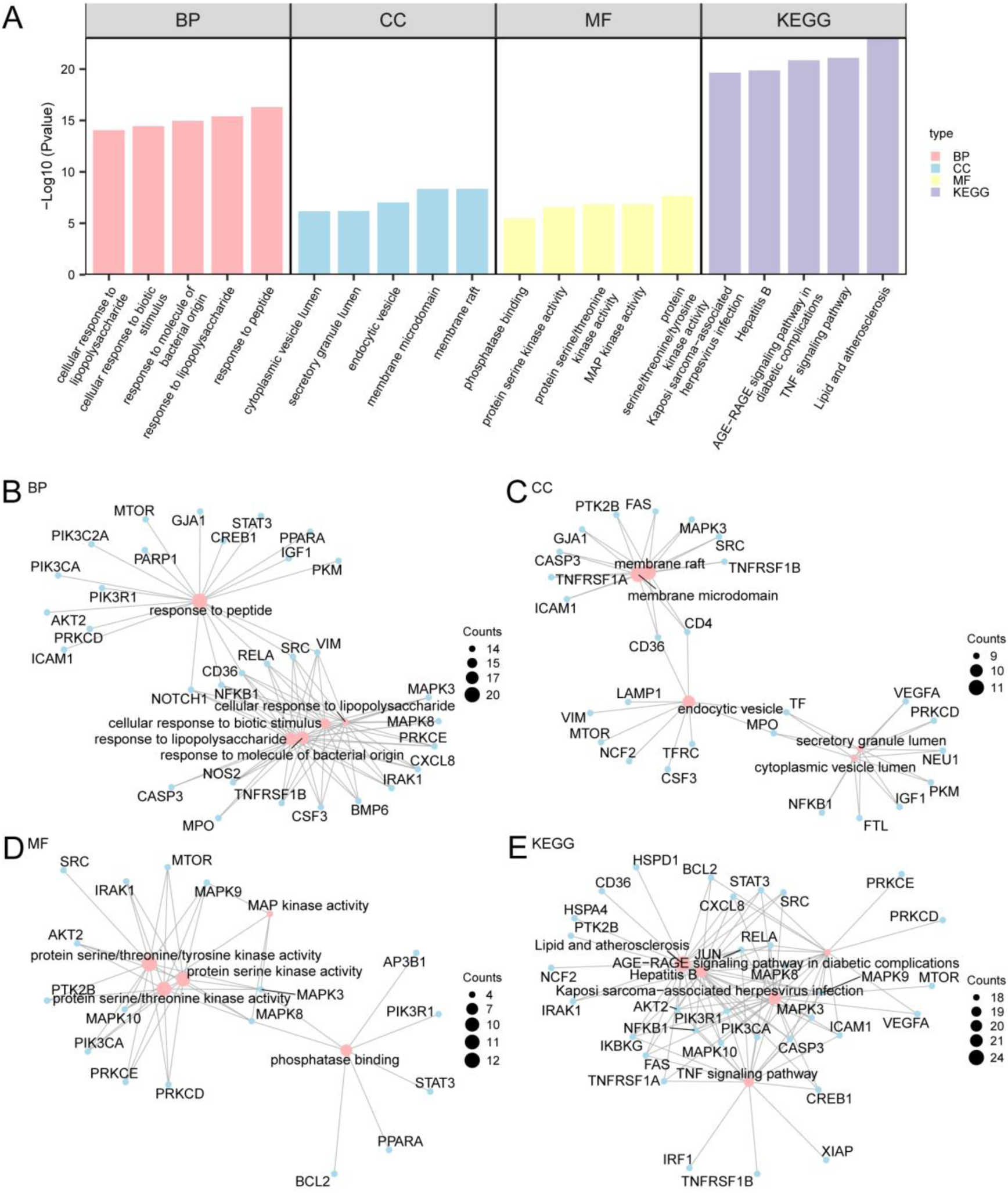
GO and KEGG Analyses of M&MARDEGs. A. GO and KEGG results of M&MARDEGs could be visualized in the bar diagram: BP, CC, MF and KEGG pathways. The abscissa coordinates are GO and KEGG terms. B-E. GO and KEGG outcomes of mitochondria & Macrophage activation-related differentially expressed genes (M&MARDEGs) can be observed from the network diagram: BP (B), CC (C), MF (D) and KEGG (E). Pink and blue nodes stand for entries and molecules, whereas lines stand for relationships of entries with molecules. M&MARDEGs, Mitochondria & Macrophage Activation-Related Differentially Expressed Genes; GO, Gene Ontology; KEGG, Kyoto Encyclopedia of Genes and Genomes; BP, Biological Process; CC, Cellular Component; MF, Molecular Function. GO and KEGG analyses were completed upon the p < 0.05 and FDR (q-value) < 0.05 thresholds, and p-value was corrected by BH.

### 3.4 GSEA

For analyzing the gene expression effect from the combined GEO Datasets on Sarcopenia incidence, GSEA was conducted based on the logFC value of each gene from Combined GEO dataset in sarcopenia versus control groups, so as to study the relation of gene expression in combined GEO datasets with BPs, CCs and MFs, and a mountain map was displayed (Fig 5A). Table 3 shows the relevant results. As observed, these genes in the Combined Datasets were closely related to Pid Notch Pathway (Fig 5B), Mebarki Hcc Progenitor Wnt Up (Fig 5C), Martinez Tp53 Targets Dn (Fig 5D), and Dasu Il6 Signaling Up (Fig 5E).

**Fig 5.**
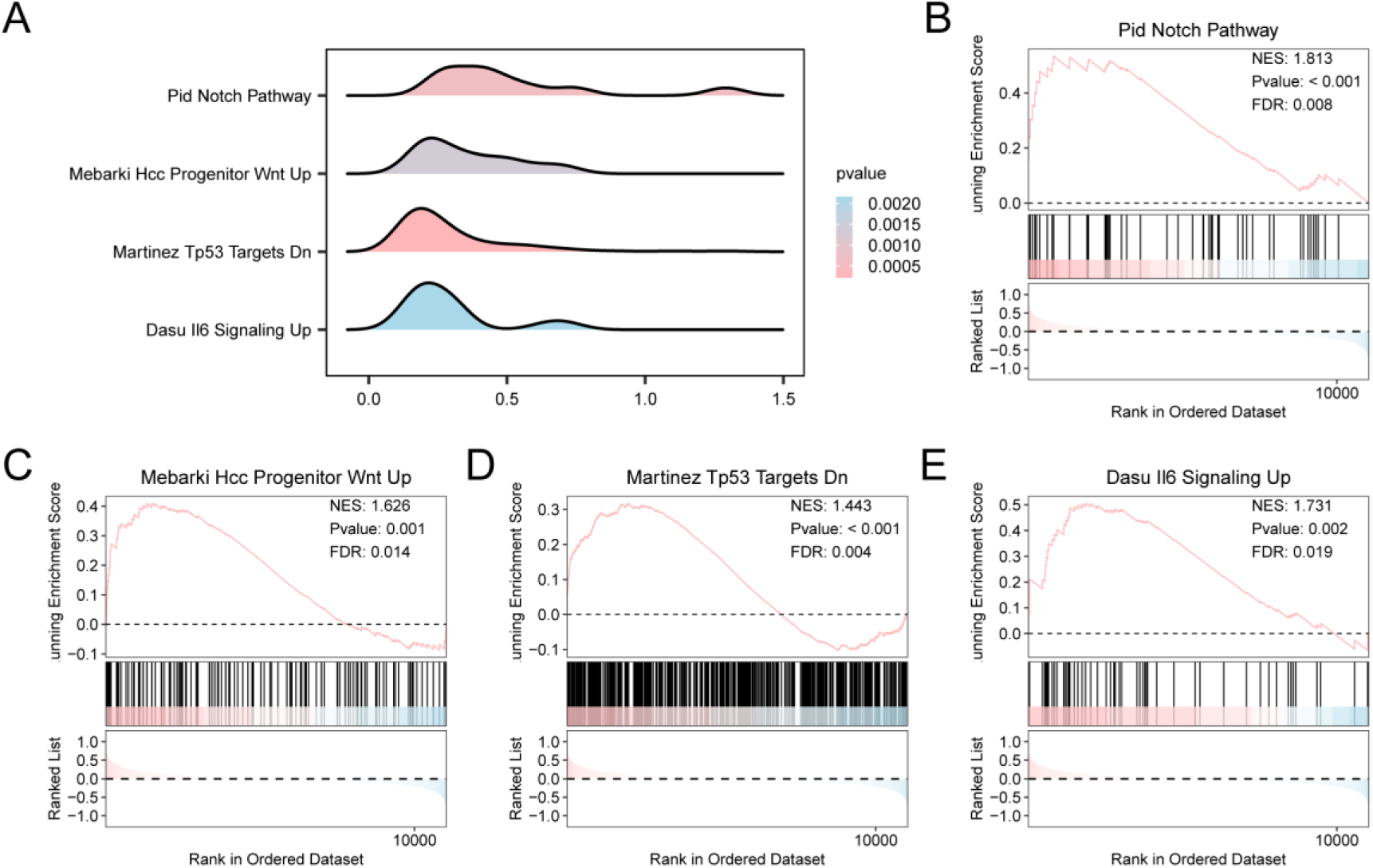
Differential Gene Expression Analysis and GSEA of Combined Datasets. A. Differential Gene Expression Analysis (GSEA) of 4 biological function mountain maps for the Combined GEO dataset. B-E. GSEA presents that the Combined Datasets was markedly related to Pid Notch Pathway (B), Mebarki Hcc Progenitor Wnt Up (C), Martinez Tp53 Targets Dn (D), Dasu Il6 Signaling Up (E). GSEA, Gene Set Enrichment Analysis. The color in mountain map indicates p-value size, with a redder color representing a lower p-value, while a bluer color standing for a greater p-value. GSEA was accomplished upon the p < 0.05 and FDR (q-value) < 0.05 thresholds, and p-value was corrected by BH.

### 3.5 Establishment of the diagnostic model for sarcopenia

First, for determining the diagnostic significance of 62 M&MARDEGs related to mitochondrial & macrophage activation in sarcopenia, this study performed the univariate logistic regression model using 62 M&MARDEGs related to mitochondrial & macrophage activation. As discovered, all these 60 M&MARDEGs showed statistical significance in this model (p < 0.05, Table S4).

Then, on the basis of 60 mitochondrial & macrophage activation-related differentially expressed genes (M&MARDEGs) via LASSO (Least Absolute Shrinkage and Selection Operator) regression was carried out for constructing the LASSO regression model, i.e. sarcopenia diagnostic model. Visualization was performed by drawing LASSO regression model diagram (Fig 6A) and LASSO variable trajectory diagram (Fig 6B). The 24 M&MARDEGs included into LASSO regression model were as follows: *IGF1, FTL, MAPK8, MAPK9, CD4, TF, CREB1, TNFRSF1B, FAS, NOS2, NOTCH1, CXCL8, VEGFA, TUG1, NFKB1, EIF4E, PIK3C2A, TFRC, VIM, BCL2, CD36, GJA1, PRKCE, LAMP1*.

**Fig 6.**
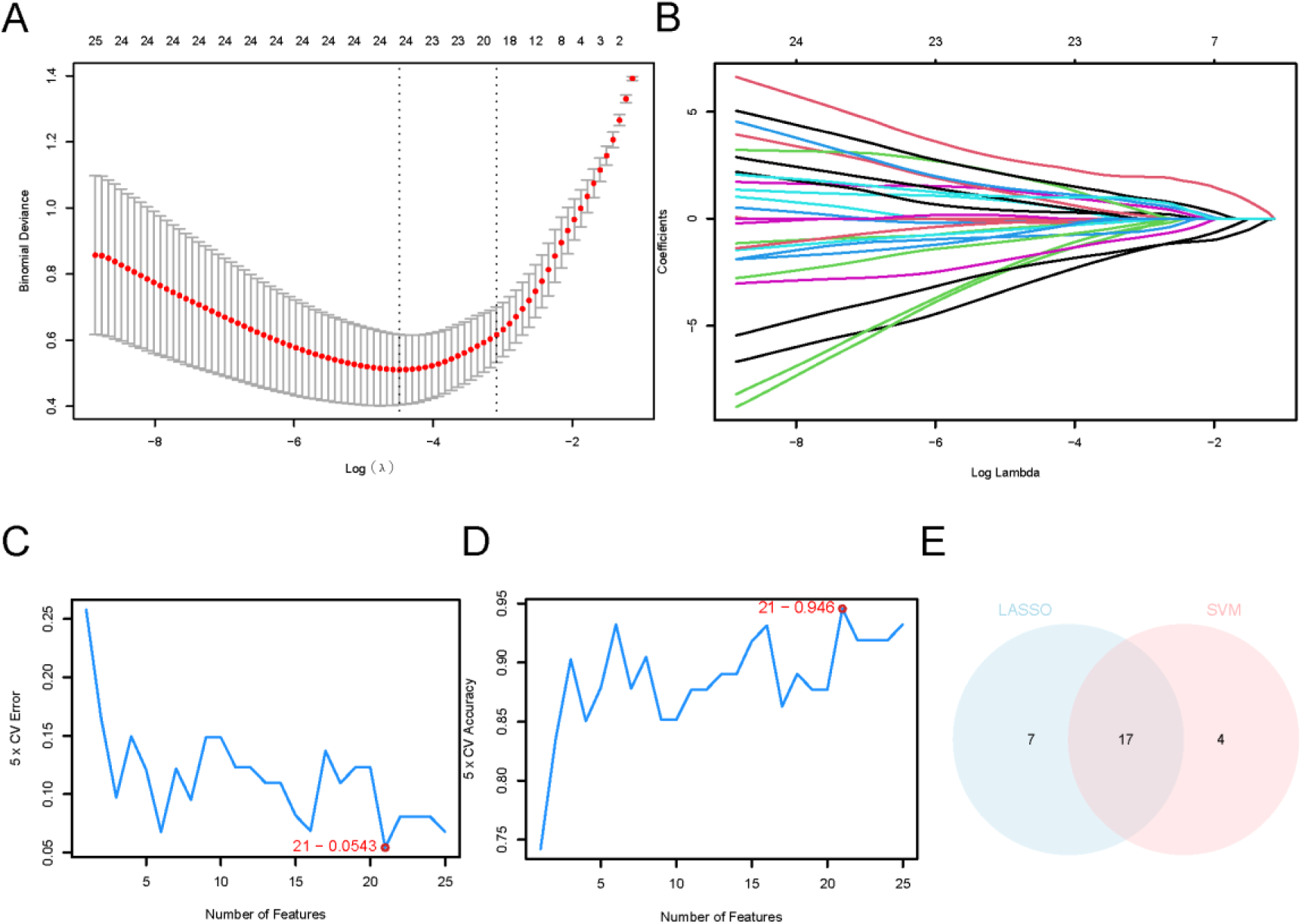
Establishment of the diagnostic model for sarcopenia. A. LASSO regression diagnostic model diagram showing mitochondria & macrophage activation-related differentially expressed genes (M&MARDEGs) in A Combined GEO dataset. B. Variable locus diagram for the LASSO diagnostic model. C. The SVM algorithm was utilized to obtain genes having the least error rate. D. The SVM algorithm was utilized to obtain genes that had the greatest accuracy. E. Venn diagram showing intersection of LASSO with SVM algorithms. LASSO: Least Absolute Shrinkage and Selection Operator. M&MARDEGs: Mitochondria & Macrophage Activation - Related Differentially Expressed Genes; SVM: Support Vector Machine.

Finally, we established a SVM model by incorporating 60 M&MARDEGs and the SVM algorithm, and later genes that had smallest error rate (Fig 6C) whereas greatest accuracy rate (Fig 6D) were acquired. As a result, the SVM model achieved greatest accuracy at a gene number of 21. The 21 M&MARDEGs related to activation of mitochondria & macrophages (M&MARDEGs) were as follows: *EIF4E, FTL, BCL2, CREB1, TF, LAMP1, TXNRD1, NFKB1, MAPK9, PKM, IGF1, CD4, VIM, TNFRSF1B, GJA1, MAPK8, TUG1, XIAP, FAS, SRC, VEGFA*.

For obtaining key genes, the M&MARDEGs in LASSO regression model were intersected with those in SVM model. A total of 17 M&MARDEGs (M&MARDEGs) were obtained for subsequent research and Fig 6E was drawn. 17 M&mardeGs (M&MARDEGs) were *IGF1, FTL, MAPK8, MAPK9, CD4, TF, CREB1, TNFRSF1B, FAS, VEGFA, TUG1, NFKB1, EIF4E, VIM, BCL2, GJA1, LAMP1*.

### 3.6 Verification of diagnostic models for sarcopenia

For validating the value of our constructed Sarcopenia diagnostic model, Nomograms based on 17 mitochondria & macrophage activation-related differentially expressed genes (M&MARDEGs) were developed to demonstrate the interrelationships in an integrated GEO dataset (Fig 7A). Altogether 9 model genes were added in multivariate regression model, which were: IGF1, FTL, MAPK8, MAPK9, TF, CREB1, FAS, VEGFA, NFKB1. We took model genes as the key genes for subsequent research. Among them, key genes FTL showed a markedly superior effect on the sarcopenia diagnostic model to additional variables.

**Fig 7.**
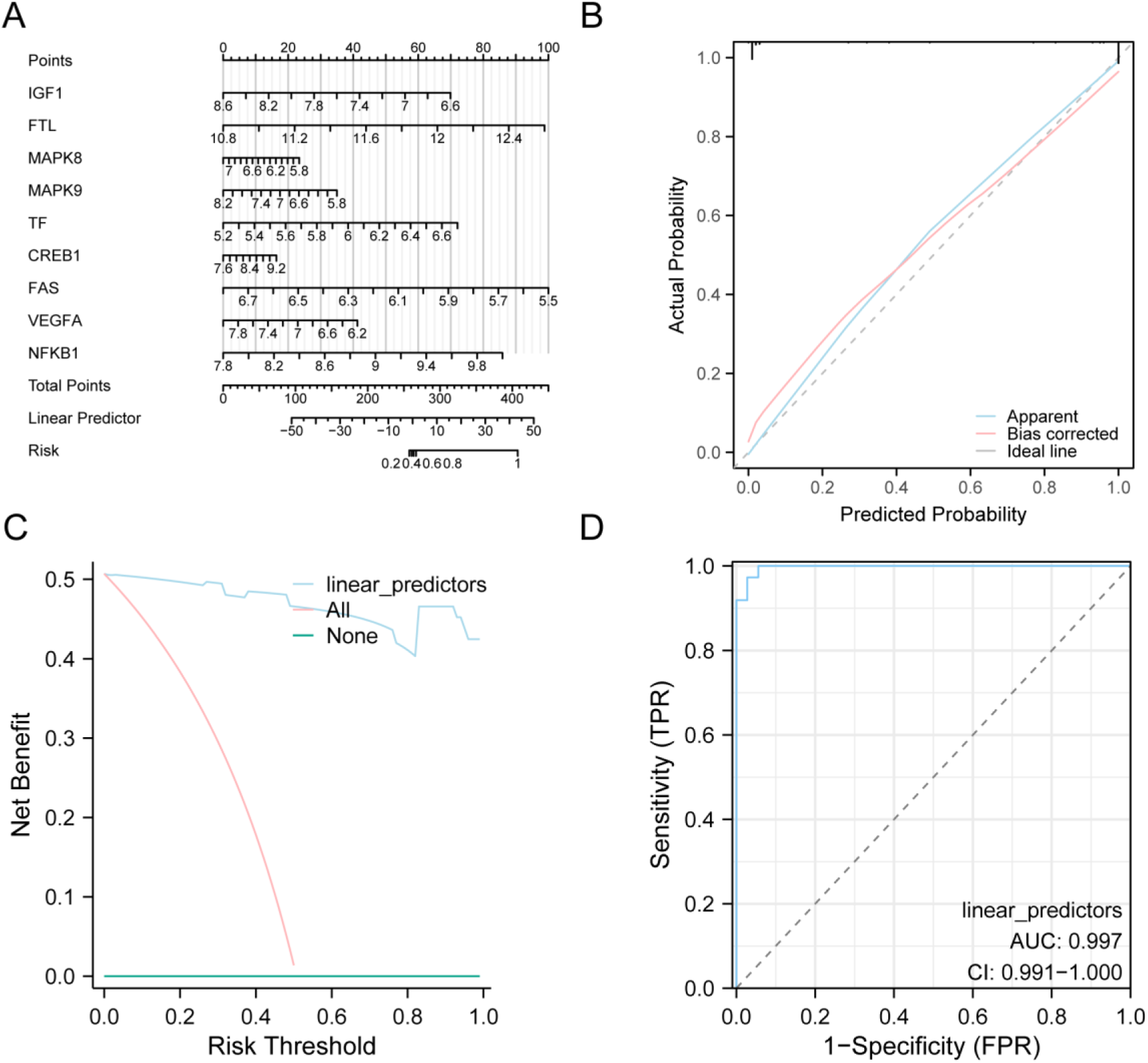
Diagnostic Performance and Verification of OA. A. Nomogram constructed by key genes from Combined GEO Datasets in a diagnostic model of Sarcopenia. B-C. Sarcopenia diagnostic model was constructed according to Calibration graph (B) and DCA graph (C) of key genes from the Combined GEO Datasets). D. Logistic regression model of Linear Predictors in ROC analysis of GEO data set (Combined Datasets). In DCA plot, the horizontal and vertical coordinate indicate Probability Threshold (or Threshold Probability) and net return separately. DCA, decision Curve analysis; ROC, Receiver Operating Characteristic; AUC has high accuracy above 0.9. AUC, Area Under the Curve.

Next, in order to evaluate the resolution and accuracy of our Sarcopenia diagnostic model, we drew the Calibration Curve through Calibration analysis. To evaluate the model prediction performance in real probability based on the fitting of model-predicted and actual probabilities in diverse situations (Fig 7B). A Calibration graph showing the sarcopenia diagnostic model suggested the slight off of the calibration line (denoted as the dashed line) from the ideal model diagonal, yet approaching match.

Key genes in the Combined Datasets evaluating the effect of our constructed sarcopenia diagnostic model on clinical utility by DCA, and findings were displayed (Fig 7C). According to our observations, the model line was stably located above the positives and negatives in the certain range, with a larger model net benefit indicating a superior model effect.

We also plotted the Receiver Operating of Linear Predictors of Logistic regression model between sarcopenia and control groups in integrated GEO Datasets Characteristic Curve (ROC) curve and result presentation (Fig 7D). The Logistic regression model of integrated GEO data set has a good diagnostic effect.

### 3.7 PPI Network

By using STRING database, PPI analysis on 9 key genes (*IGF1, FTL, MAPK8, MAPK9, TF, CREB1, FAS, VEGFA, NFKB1*) was conducted. Under the minimal required interaction score > 0.150 (minimal required interaction score: low confidence (0.150), we drew the PPI network diagram (Fig 8A). Similar Genes of 9 key genes (*IGF1, FTL, MAPK8, MAPK9, TF, CREB1, FAS, VEGFA, NFKB1*) were also predicted at GeneMANIA website, followed by mapping of this PPI network to observe the physical interaction relationship between them and the shared protein domain. Finally, similar genes of 9 key genes (*IGF1, FTL, MAPK8, MAPK9, TF, CREB1, FAS, VEGFA, NFKB1*) were predicted at GeneMANIA website and the interaction network was constructed for observing the shared protein domains, physical interactions, gene-gene interactions, and other relevant information among them (Fig 8B).

**Fig 8.**
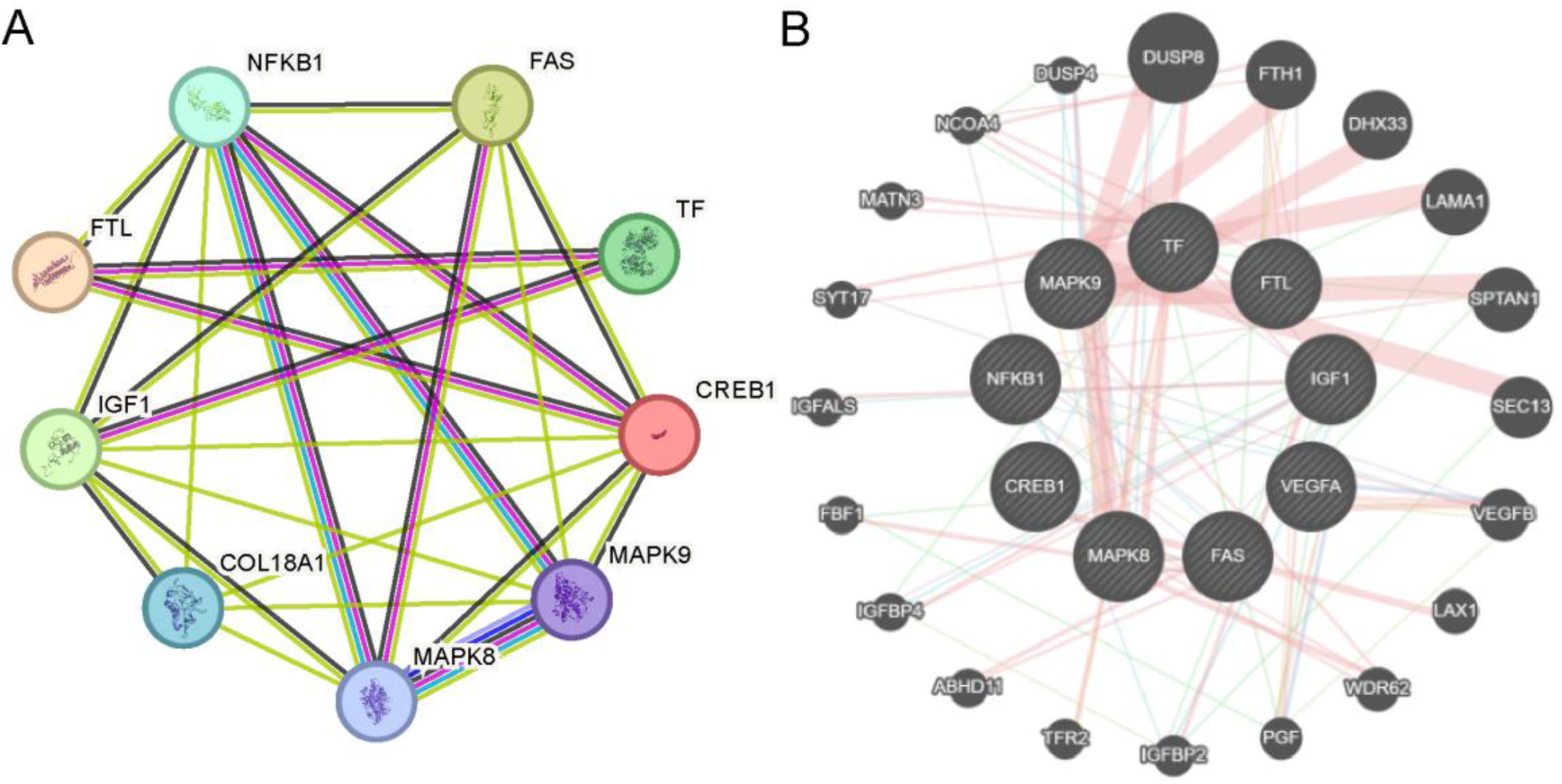
PPI Network. A. The PPI Network for key genes. B. key genes predicting the PPI network of functionally similar genes. Genes are represented by circular nodes, whose size is related to gene characteristics and attributes. Lines stand for between-gene relations, interactions and functional connections, with lines thick and thin representing strong associations or significant interactions. PPI Network: Protein-Protein Interaction Network.

### 3.8 Establishment of regulatory network

First, key genes-related miRNAs were acquired by TarBase database (StarBase database). Thereafter, Cytoscape was applied in visualizing our created mRNA-miRNA regulatory network (Fig 9A). In this network, 8 key genes and 172 miRNAs were included. For details, see Table S5.

**Fig 9.**
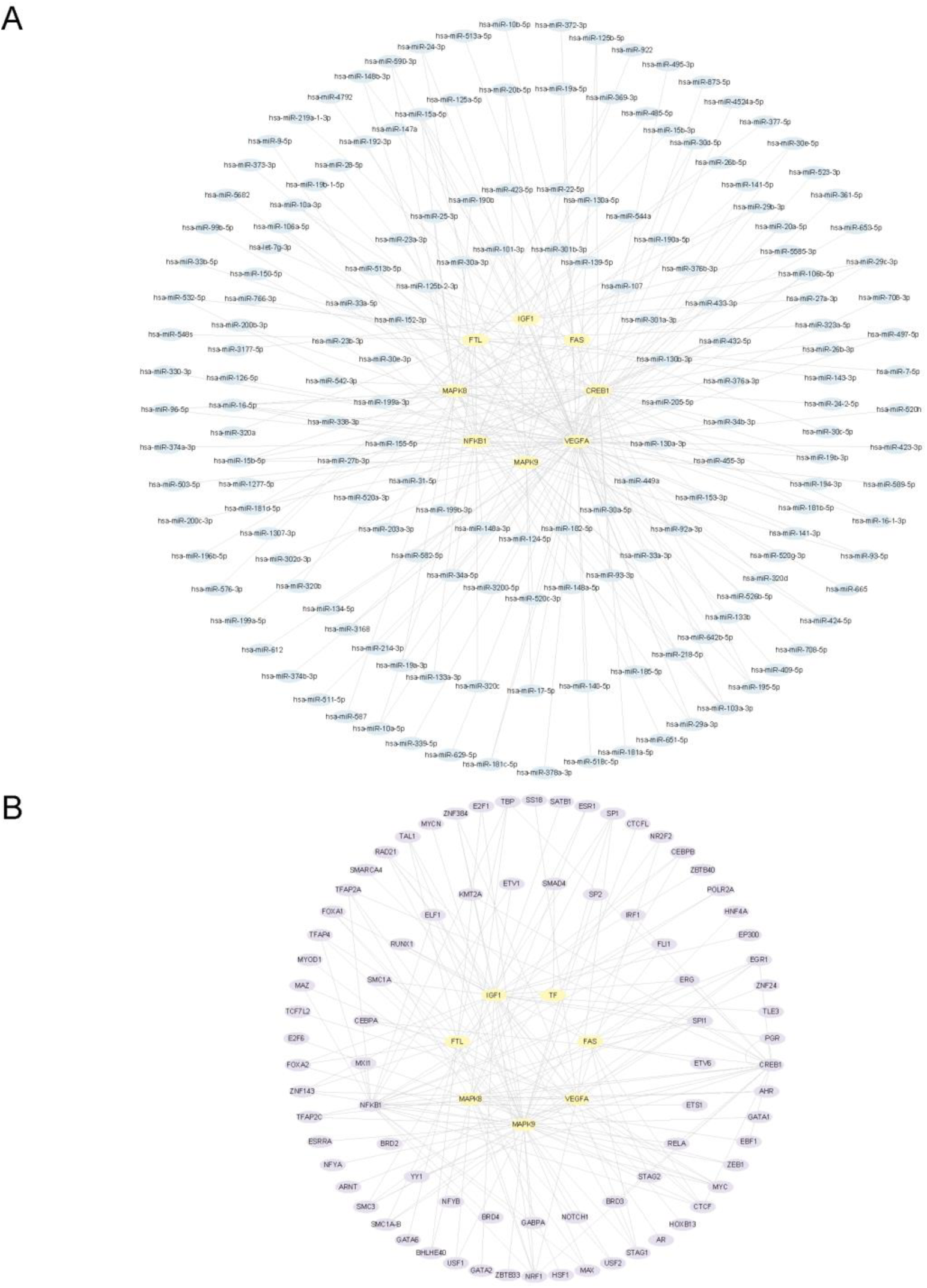
The key gene Regulatory Network. The mRNA-miRNA (A) and mRNA-TF (B) regulatory networks for key genes. TF, Transcription Factor; Yellow for mRNA, blue for miRNA, and purple for TF.

Then, key gene-binding TFs were collected through ChIPBase database, followed by mRNA-TF regulatory network construction and visualization with Cytoscape (Fig 9B). Of these, there were 9 key genes and 81 TFs included. Table S6 shows more details.

### 3.9 Differential expression of key genes in diverse groups

There were 9 key genes (IGF1, FTL, MAPK8, MAPK9, TF, CREB1, FAS, VEGFA) shown in Fig 10A by group comparison. The NFKB1 differential expression in control and sarcopenia groups was analyzed in a Combined GEO Datasets. As discovered, five key genes (IGF1, FTL, MAPK9, CREB1, NFKB1) showed significantly different expression levels in Control versus Sarcopenia groups from the integrated GEO datasets (p < 0.001). Besides, key genes (MAPK8, TF, VEGFA) had significantly different expression levels in Control versus Sarcopenia groups from the Combined Datasets (p < 0.01). Similarly, FAS also displayed significantly different expression in Control versus Sarcopenia groups from the Combined Datasets (p < 0.05).

**Fig 10.**
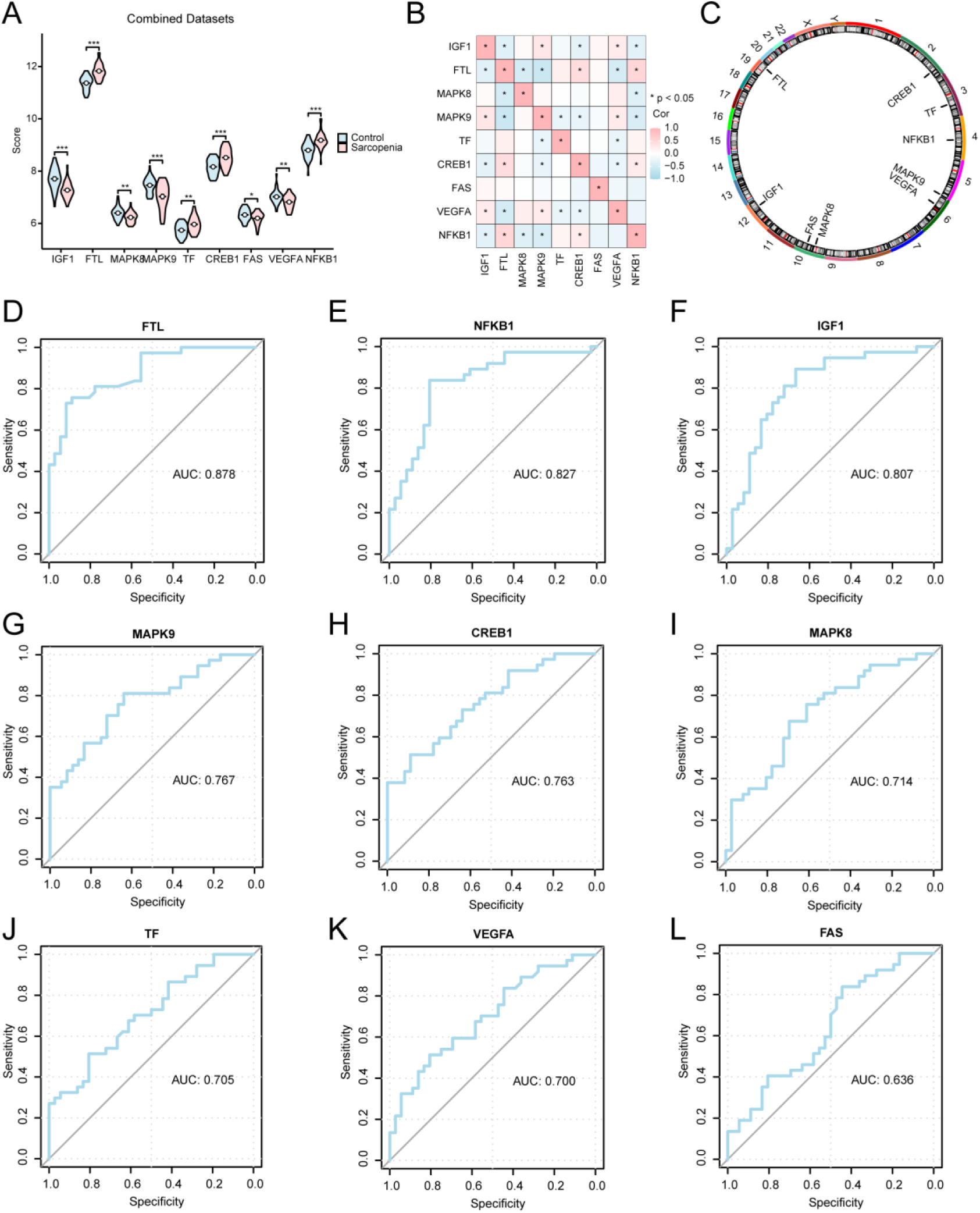
Analysis of differential Key Gene expression among diverse subgroups in Combined GEO Datasets. A. Grouping comparison in Control versus sarcopenia groups in the Combined GEO Datasets. B. Correlation analysis among key genes. C. Chromosomal mapping of key genes in human body. D-L. Key genes FTL (D), NFKB1 (E), IGF1 (F), MAPK9 (G), CREB1 (H), MAPK8 (I), TF (J), VEGFA (K), FAS (L) ROC curve analysis for the Combined Datasets. ***P < 0.001 and **P < 0.01 stand for highly statistical significance, while *P < 0.05 represents statistical significance. An AUC value approaching 1 represents superior diagnostic effect. The AUCs of 0.7-0.9 and 0.5-0.7 show certain and low accuracy, separately. ROC: Receiver Operating Characteristic. AUC: Area Under Curve. Pink and blue stand for sarcopenia and control groups.

Based on 9 key gene levels (IGF1, FTL, MAPK8, MAPK9, TF, CREB1, FAS, VEGFA, NFKB1) in Combined Datasets complete expression matrix for correlation analysis and mapping of correlation heat map (Fig 10B). As a result, key genes such as NFKB1 and FTL were positively correlated, whereas MAPK9 and FTL were negatively correlated.

We also annotated the positions of 9 key genes (IGF1, FTL, MAPK8, MAPK9, TF, CREB1, FAS, VEGFA, NFKB1) in human chromosomes and visualized them in a circular graph (Fig 10C). As shown in the figure, key genes CREB1, TF, NFKB1, MAPK9, VEGFA, MAPK8 and FAS, IGF1 and FTL are on chromosomes 2, 3, 4, 5, 6, 10 and 12, respectively.

Finally, map 9 key genes (IGF1, FTL, MAPK8, MAPK9, TF, CREB1, FAS, VEGFA, etc.).

NFKB1) in the Combined Datasets. ROC analysis results are displayed (Fig 10D-L). Differential expression of key genes (IGF1, FTL, MAPK8, MAPK9, TF, CREB1, VEGFA, NFKB1) in the Combined Datasets exhibited some accuracy (0.7 < AUC < 0.9) across diverse groups. FAS expression in the Combined Datasets had lower accuracy across subgroups (0.5 < AUC < 0.7).

ROC curves of 9 Key Genes (IGF1, FTL, MAPK8, MAPK9, TF, CREB1, FAS, VEGFA, NFKB1) were plotted in dataset GSE136344, and results of AUC <0.6 were not displayed (Fig 11A-E). The differential Key Gene expression (FTL and FAS) in GSE136344 dataset exhibited a certain accuracy across diverse groups (0.7 < AUC < 0.9). Differences in expression of Key Genes (MAPK9, VEGFA and CREB1) in the dataset GSE136344 showed lower accuracy across subgroups (0.5 < AUC < 0.7).

**Fig 11.**
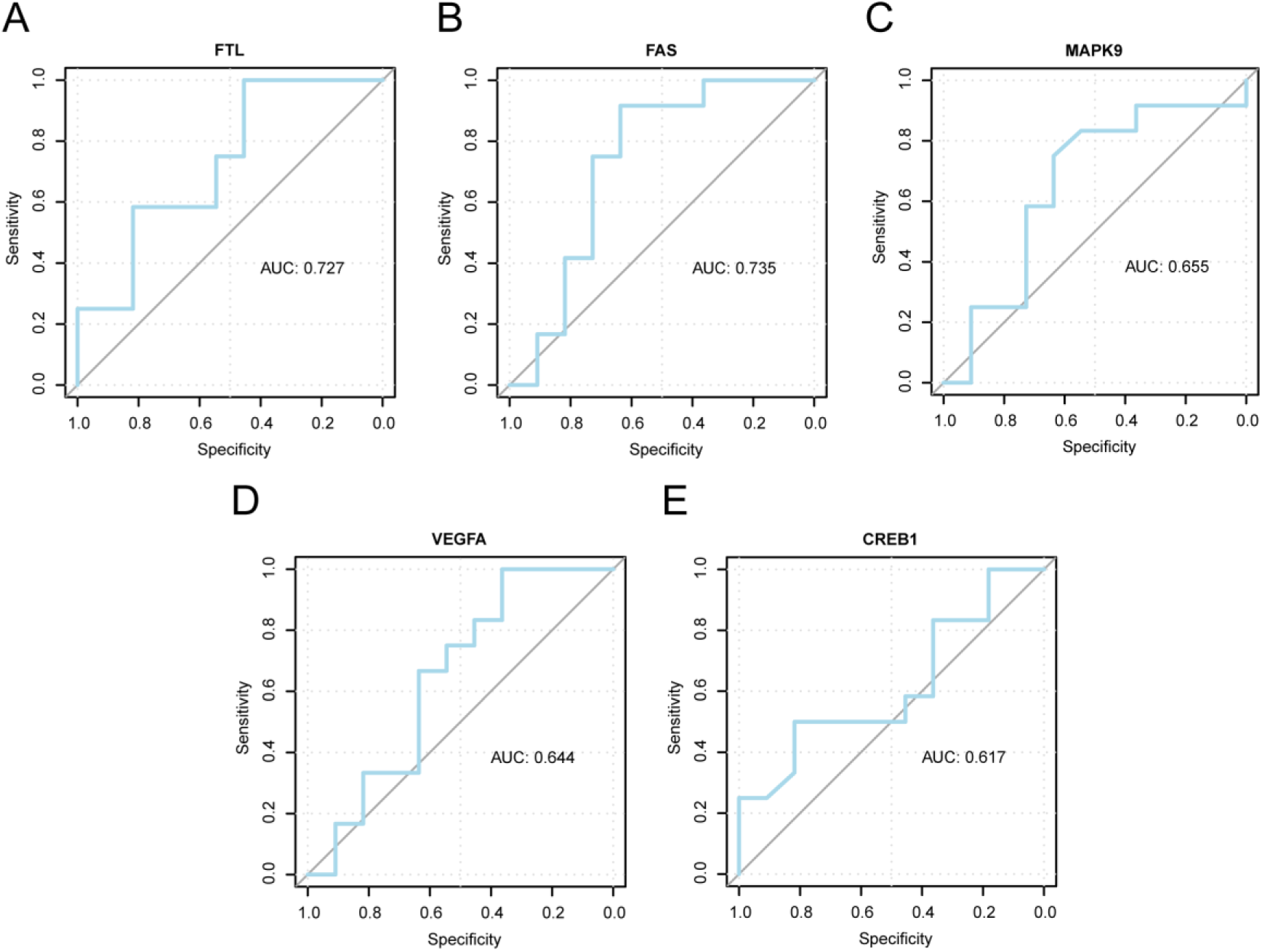
Analysis of expression differences of Key Genes among diverse subgroups in dataset GSE136344. A-E. ROC curve analysis for key genes FTL (A), FAS (B), MAPK9 (C), VEGFA (D), CREB1 (E) in dataset GSE136344. An AUC approach 1 in ROC curve suggests the better diagnostic performance. The AUCs of 0.7-0.9 and 0.5-0.7 showed certain and low accuracy separately. ROC: Receiver Operating Characteristic. AUC: Area Under Curve.

### 3.10 Immunoinfiltration analysis

Based on 9 key genes (IGF1, FTL, MAPK8, MAPK9, TF, CREB1, FAS, VEGFA) we calculated the Mitochondria & Macrophage Activation Score (M&MA. Score) of each sample according to the NFKB1 expression matrix in a Combined Datasets (GEO). In addition, median expression value of the mitochondria & macrophage activation score (M&MA.Score) classified sarcopenia patients in high score (HighScore) or low score (LowScore) group. For the Combined GEO dataset, its expression matrix was used for calculating 28 immune cell infiltration levels in sarcopenia sample using ssGSEA algorithm.

First, the differential immune cell infiltration levels in diverse groups were demonstrated through grouping comparison plots. Fig 12A showed that Central memory CD8 T+ cells and Effector memory CD8 T+ cells reached the significance level (p < 0.05).

**Fig 12.**
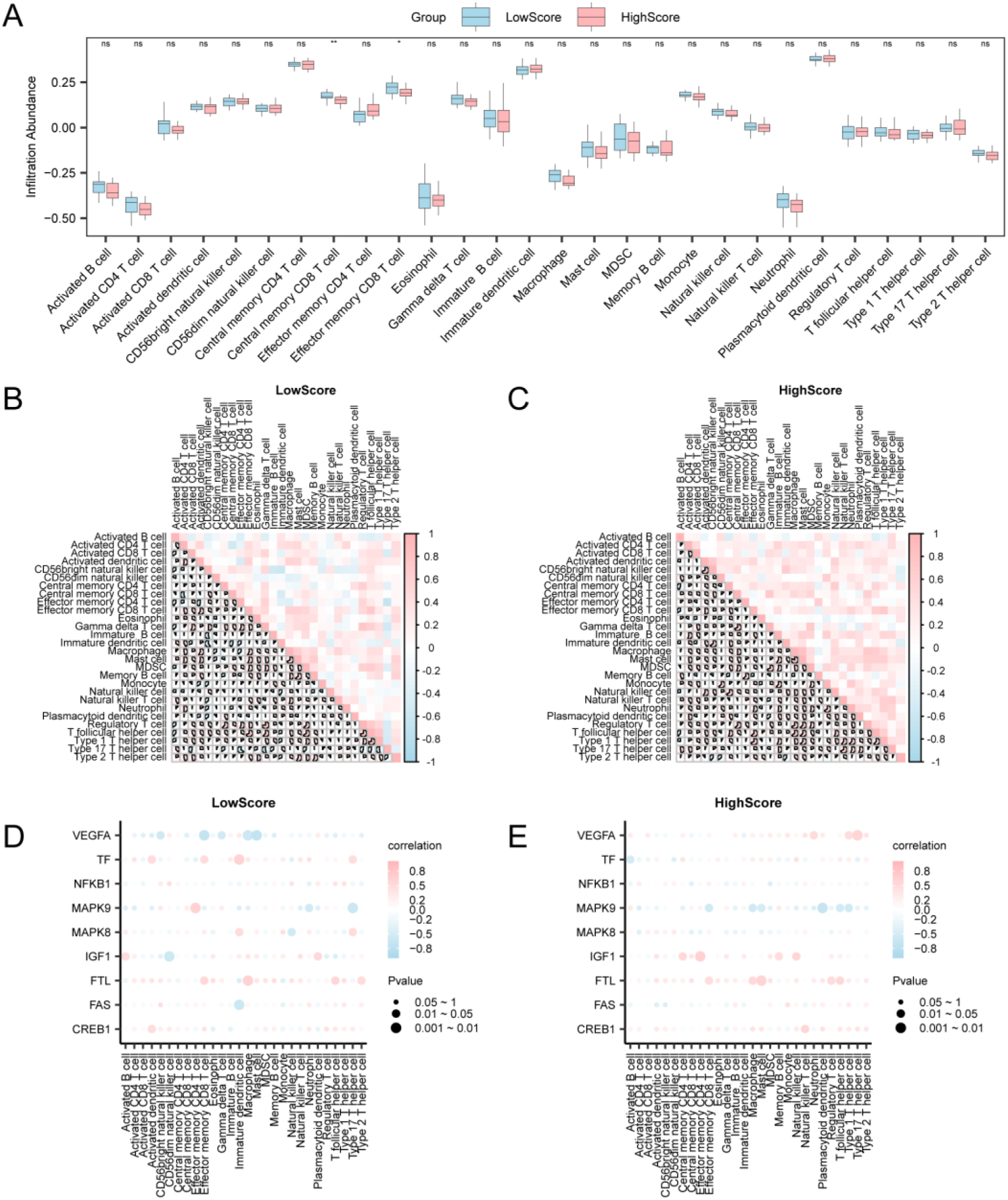
Risk Group Immune Infiltration Analysis with ssGSEA Algorithm. A. Immune cells Mitochondria & Macrophage Activation in LowScore group of sarcopenia samples and Mitochondria & Macrophage A grouping comparison diagram showing activation in HighScore group. Mitochondria & Macrophage Activation LowScore group (B) and Mitochondria & macrophage activation in the B-C. Sarcopenia sample Results of correlation analysis on the abundance of immune cell infiltration in Macrophage Activation group (C) with HighScore. D-E. Correlation bubble plots showing the relation of immune cell infiltration levels with mitochondria & macrophage activation-related differentially expressed genes (M&MARDEGs) in low-score (LowScore) (D) and high-score (HighScore) (E) groups of sarcopenia samples. ssGSEA, single-sample Gene-Set Enrichment Analysis. Ns, not significant (p ≥ 0.05); *p < 0.05, statistical significance; **p < 0.01, highly statistical significance. The |r value (correlation coefficient)| < 0.3, 0.3-0.5, 0.5-0.8 and > 0.8 represents weak/no, weak, moderate and strong correlation separately. Mitochondria & Macrophage Activation group with LowScore is in blue, And pink for Mitochondria & Macrophage Activation high score group (HighScore). Pink and blue stand for positive and negative correlation separately, with color depth indicating the correlation strength.

Later, a correlation heatmap was drawn to present correlations of 28 immune cell infiltration levels in sarcopenia samples (Fig 12B-C). As observed, many immune cells Mitochondria & Macrophage Activation LowScore group in the sarcopenia sample showed strong correlation. Gamma delta T cell and Regulatory T cell were most positively correlated (r = 0.75, p < 0.05) (Fig 12B). Mitochondria & Macrophage Activation group with HighScore showed strong positive correlation with most immune cells. Macrophage and Mast cell were most positively correlated (r = 0.812, p < 0.05) (Fig 12C).

Eventually, we showed the associations of M&MARDEGs with immune cell infiltration levels through plotting the correlation bubble diagram (Fig 12D-E). As discovered, many immune cells in sarcopenia sample showed strong correlation with Mitochondria & Macrophage Activation group with LowScore, VEGFA gene and mast cell were most negatively correlated (r = -0.707, p < 0.05) (Fig 12D). Mitochondria & Macrophage Activation group with HighScore showed strong correlation with most immune cells. The MAPK9 gene and Plasmacytoid dendritic cell were most negatively correlated (r = -0.661, p < 0.05) (Fig 12E).

### 3.11 Establishment of sarcopenia subtypes

To explore disease subtypes among sarcopenia samples from the sarcopenia dataset, the R package ConsensusClusterPlus was used, based on nine Key Genes (IGF1, FTL, MAPK8, MAPK9, TF, CREB1, FAS, VEGFA, etc.). The expression of NFKB1 in the Sarcopenia sample in the Sarcopenia dataset was utilized for identifying the diverse disease subtypes of Sarcopenia sample using consistent cluster analysis, and finally 2 Sarcopenia subtypes were identified (Fig 13A-C): Subtype A (Cluster1, containing 18 samples) and subtype B (Cluster2 containing 19 samples). PCA plots revealed significantly different NFKB1 expression between both disease subtypes (Fig 13D).

**Fig 13.**
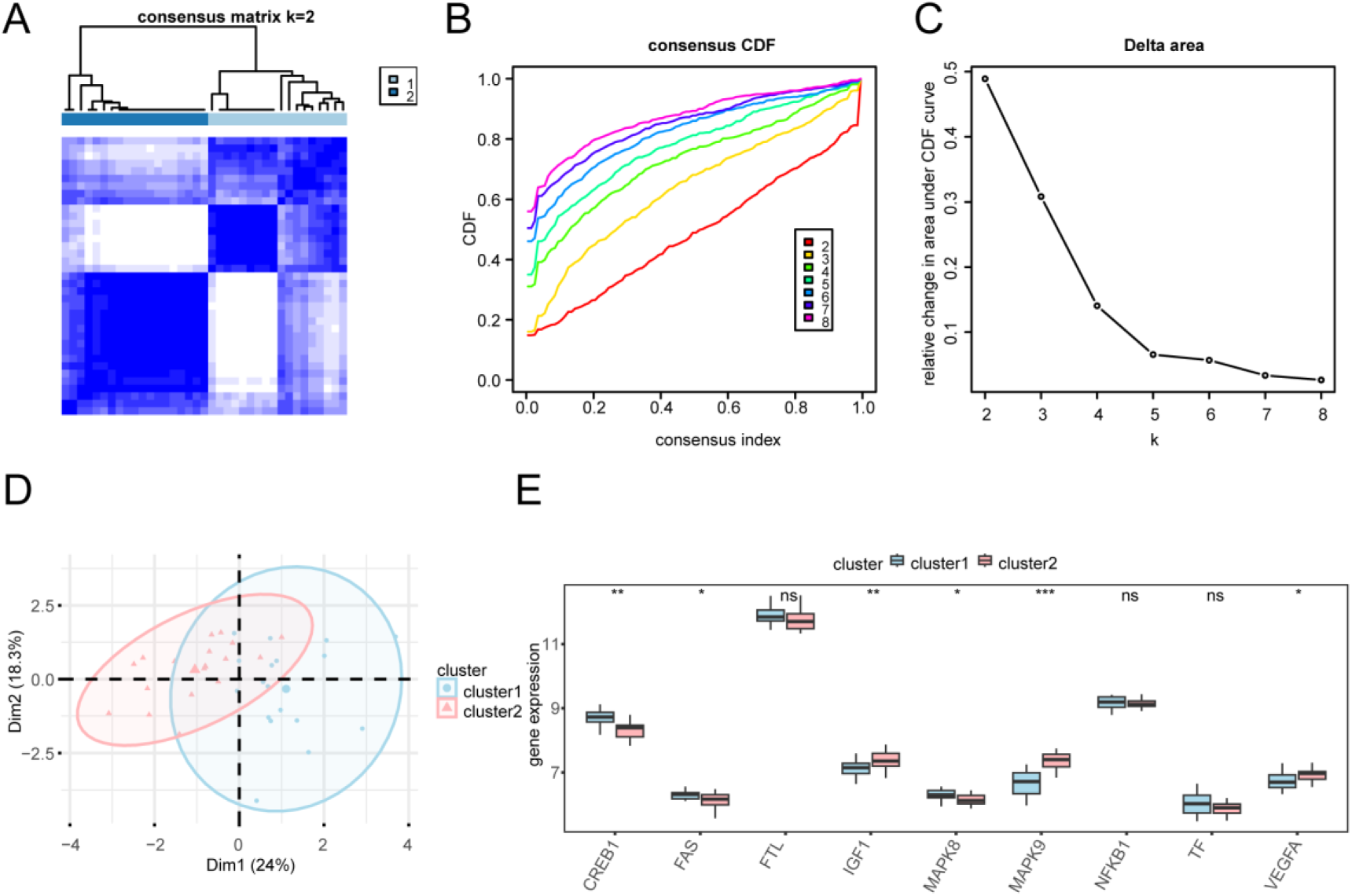
Consensus Clustering Analysis of Sarcopenia. A. Consistent clustering of a sarcopenia sample from the sarcopenia dataset. B-C. Consistent CDF graph (B) and Delta graph (C) for consistent clustering. D. Sarcopenia, PCA diagram showing 2 disease subtypes. E. Subgroup comparison diagram for key genes between two sarcopenia subtypes. CDF, Empirical Cumulative Distribution Function; And PCA, Principal Component Analysis. ***P < 0.001 and **P < 0.01 stand for highly statistical significance; *P < 0.05 represents statistical significance; ns is not significant (P > 0.05). Blue is subtype A (Cluster1) and pink is subtype B (Cluster2).

Finally, in order to better validate differential key gene expression among sarcopenia subtypes, our analysis results of key gene expression with two sarcopenia subtypes were presented in the group comparison diagram (Fig 13E). Fig 13E showed significantly different key gene MAPK9 level between two disease subtypes (p < 0.001). Additionally, key genes (CREB1 and IGF1) showed highly significant difference between two disease subtypes (p < 0.01). Key genes (FAS, MAPK8 and VEGFA) exhibited significantly different levels between two disease subtypes (p < 0.05).

### 3.12 Immunoinfiltration analysis

For the integrated GEO dataset, its expression matrix was utilized for calculating 28 immune cell infiltration levels in the Sarcopenia sample with ssGSEA algorithm. First, the differential immune cell infiltration levels of diverse groups were demonstrated through grouping comparison plots. Fig 14A showed that Effector memory CD4 T cell, Effector memory CD8 T cell, and Macrophage reached significant differences (p < 0.05).

**Fig 14.**
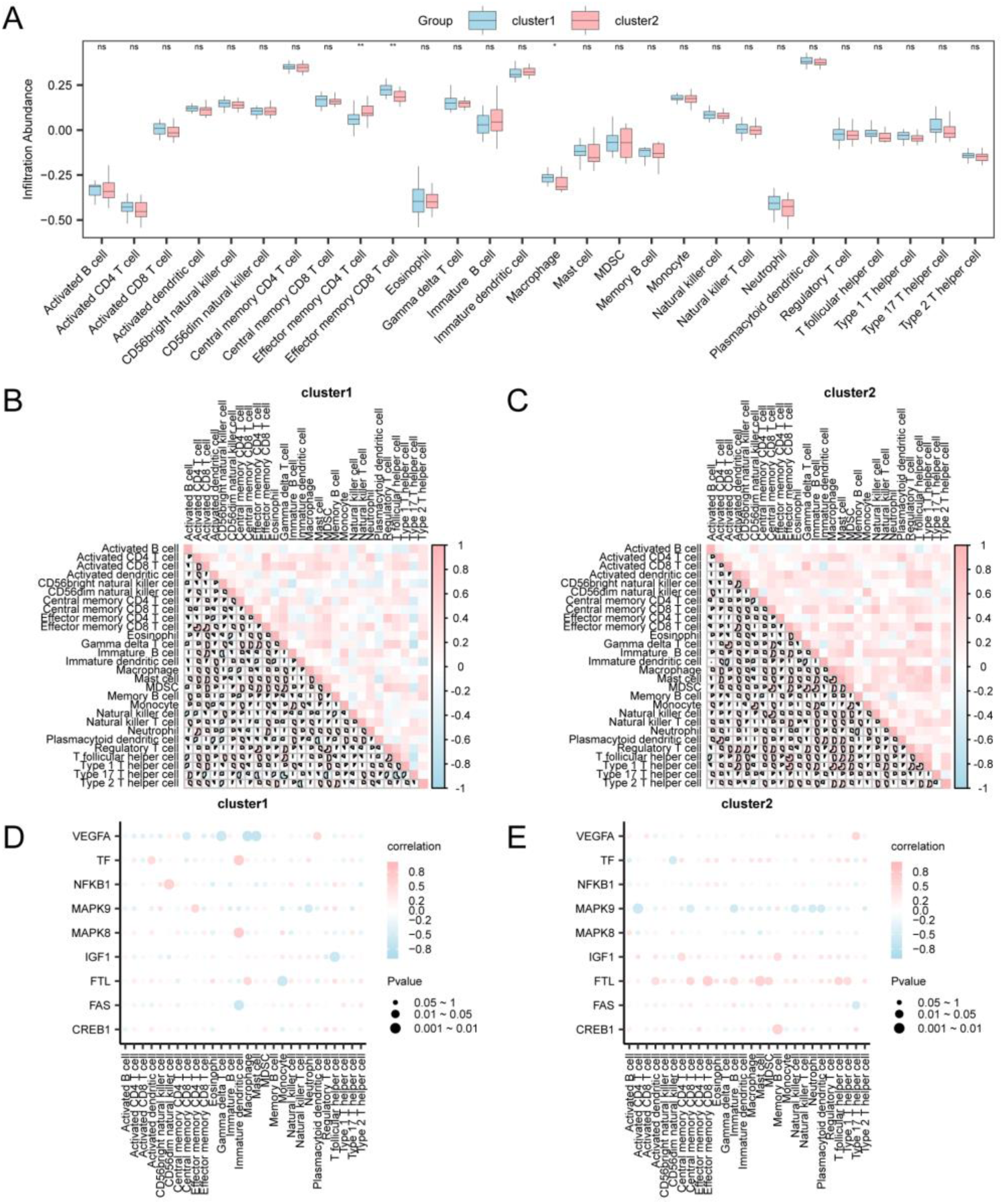
Risk Group Immune Infiltration Analysis with ssGSEA Algorithm. A. Grouping comparison diagram for immune cells of subtype (cluster1) versus subtype (cluster2) samples of Sarcopenia. B-C. Correlations of immune cell infiltration levels between subtype (cluster1) (B) and subtype (cluster2) (C) groups in Sarcopenia samples presented. The immune cell infiltration levels in subtype (cluster1) (D) versus subtype (cluster2) (E) groups. D-E. Sarcopenia sample correlated with M&MARDEGs related to activation. ssGSEA, single-sample Gene-Set Enrichment Analysis. ***P < 0.001 and **P < 0.01 stand for highly statistical significance; *P < 0.05 represents statistical significance; ns is not significant (P > 0.05). The blue group is subtype (cluster1) and the pink group is subtype (cluster2). Pink is positively correlated, blue is negatively correlated, with color depth indicating correlation strength.

Subsequently, we drew a correlation heatmap for presenting correlations of 28 immune cell infiltration levels in sarcopenia samples (Fig 14B-C). As discovered, many immune cells of sarcopenia subtype (cluster1) group had potent correlation. The Activated CD4 T cell and Effector memory CD8 T cell were most positively correlated (r = 0.759, p < 0.05) (Fig 14B). Many immune cells of subtype (cluster2) group were strongly and positively correlated, meanwhile, both Effector memory CD8 T cell and T helper cell showed significant positive correlation (r = 0.802, p < 0.05) (Fig 14C).

Eventually, we showed the associations of M&MARDEGs with immune cell infiltration levels through correlation bubble diagram (Fig 14D-E). As observed, many immune cells of sarcopenia subtype (cluster1) group were strongly correlated, among which, MAPK8 was most positively correlated with immune Immature dendritic cell (r = 0.688, p < 0.05) (Fig 14D); many immune cells of subtype (cluster2) group were strongly correlated, and the gene FTL was most positively correlated with immune cell Effector memory CD8 T cell (r = 0.634, p < 0.05) (Fig 14E).

## Discussion

Sarcopenia, also known as skeletal muscle loss, is an important health issue that is receiving increasing attention globally. In 2016, sarcopenia was officially classified by the World Health Organization (WHO) as a stand-alone disease, marking international recognition of its importance and underscoring a need for more in-depth research and development of effective coping strategies for the condition. Sarcopenia is featured by the progressive skeletal muscle mass loss, functional and strength decline. Sarcopenia impairs the life quality among older adults, and induces an elevated risk of complications like fracture and fall, which can lead to incapacitation or even death in severe cases [39]. Unfortunately, there is still a lack of standardized criteria and efficient diagnostic tools for diagnosing sarcopenia.

In terms of treatment, effective measures for sarcopenia are similarly limited. Although exercise training, especially resistance training, is considered one of the key tools to improve muscle condition, the development of individualized and easy-to-implement exercise programs needs to be further explored [40–41]. Meanwhile, nutritional supplementation such as increased protein intake is also thought to be helpful in maintaining and restoring muscle mass, but there is no consensus on the specific optimal dosage and combination. Thus, the prevention and treatment of sarcopenia face many difficulties and urgently require the joint efforts of the medical community and the society, with a view to finding more effective ways to combat this widespread health problem.

In the comprehensive bioinformatics analyses we conducted, we took a systematic and comprehensive approach to explore and identify potential molecular markers that are critical for understanding complex biological processes and developing new diagnostic or therapeutic strategies. The present work successfully obtained 62 DEGs that are closely related to mitochondrial function and macrophage activation. These genes showed significant expression changes, possibly reflecting their roles in mitochondrial energy metabolism, immune response regulation, and inflammatory processes. This finding not only contributes to further understanding the disease pathogenic mechanisms like sarcopenia, but also provides valuable candidate targets for future research.

Based on the above gene expression data and other relevant biological information, our team also established and rigorously verified the new diagnostic model for sarcopenia. This model combines multi-dimensional data analysis and aims to improve the sensitivity and specificity of early detection of sarcopenia, which thus provides an accurate and reliable diagnostic tool for clinicians. The PKM gene encodes the muscle-type isoform of pyruvate kinase, an important enzyme responsible for catalyzing the last step in glycolysis [42]. PKM is selectively spliced to produce two major protein isoforms, PKM1 and PKM2. In a healthy state, PKM in skeletal muscle exists primarily as PKM1, ensuring efficient glycolytic processes to support the rapid energy supply required for muscle contraction [43–44]. PKM1 is essential for maintaining energy homeostasis in muscle because it rapidly converts glucose to ATP, which provides the necessary fuel for high-intensity exercise. These researches will help reveal more secrets about skeletal muscle health and disease, and bring new hope for the treatment of related disorders. In this study, we systematically identified the BPs, MFs and CCs enriched by DEGs, as well as the major metabolic and signaling pathways in which they are involved through GO and KEGG pathway analyses. Such analyses not only provide theoretical support for our experimental data, but also lay a basis for exploring the underlying molecular mechanisms.

The “TNF pathway” is a core regulatory mechanism of the inflammatory response. It is a key component of intercellular communication, mediated mainly by TNF and its receptors (TNFRs), and regulates a series of BPs, such as inflammatory response, apoptosis, immune response, and cell survival [45–46]. Upon activation of TNFR1, other molecules such as RIP1 (receptor-interacting protein 1) and TRAF2 (TNF receptor-associated factor 2), generating a complex. This generated complex can further activate the IKK (IkB kinase) complex, resulting in IkB degradation and phosphorylation, releasing NF-kB, making its way to cell nucleus to initiate target gene transcription. These genes products are mostly pro-inflammatory factors like IL-1, IL-6, and COX-2, which promote the onset and development of the inflammatory response [47]. According to a previous study, TNF administration apparently increased the glutamate level, causing stress to astrocytes in the process of neuroinflammation [48]. Therefore, this study is consistent with previous research findings and TNF pathway has a key effect on mitochondrion after sarcopenia.

The enrichment of metabolic pathways including “glycolysis/glycolysis”, “fatty acid metabolism” and “amino acid metabolism” reveals energy metabolic reprogramming in cells in response to different stimuli. In particular, under hypoxia or nutrient deprivation, cells may preferentially choose the fast but less efficient glycolytic pathway to meet energy demands, even under normal mitochondrial function. In addition, changes in fatty acid and amino acid metabolism may reflect adjustments in cellular preferences for substrate utilization in response to different metabolic stresses.

The novel sarcopenia diagnostic model, constructed via LASSO regression and SVM algorithms with 17 pivotal genes (e.g., *IGF1, FTL, MAPK8*), demonstrated robust predictive accuracy, highlighting its potential for early clinical screening. Functional enrichment analysis confirmed the biological relevance of these genes in sarcopenia-related pathways, including muscle atrophy and inflammation. Future efforts should focus on external validation in diverse cohorts, integration with multimodal biomarkers, and exploration of therapeutic response prediction to advance personalized management.

This study successfully constructed a PPI network involving nine key genes, systematically revealing the complex synergistic relationships among sarcopenia-associated proteins. Core hub nodes within the network (e.g., *MAPK8, VEGFA, and CREB1*) play pivotal roles in regulating inflammatory responses, apoptosis, and muscle metabolism, suggesting that they may mediate abnormal activation of downstream signaling pathways (e.g., MAPK/ERK and VEGF signaling pathways) through the formation of dynamic complexes, thereby driving sarcopenia progression. This discovery not only provides novel insights into the molecular mechanisms of sarcopenia but also highlights potential therapeutic targets such as MAPK8 and VEGFA. However, the biological significance of the PPI network requires further validation. Physical Interaction Validation: Co-immunoprecipitation (Co-IP) or fluorescence resonance energy transfer (FRET) assays are needed to confirm the physical interactions of specific complexes (e.g., MAPK8-CREB1) and their regulatory effects on downstream pathways (e.g., myocyte differentiation or autophagy). Functional CRISPR Validation: CRISPR/Cas9-based knockout or overexpression experiments targeting key hub genes (e.g., VEGFA) could verify their causal roles in muscle atrophy and the functional necessity of their interactions within the network. Dynamic Network Analysis: Integration of single-cell transcriptomic or time-series data would help elucidate dynamic changes in the PPI network across disease stages or specific cell types (e.g., muscle satellite cells), ultimately revealing clinically actionable intervention windows. This PPI analysis establishes a framework for understanding sarcopenia mechanisms, but its translational potential hinges on multidisciplinary validation and multi-omics integration.

Through consensus clustering, we obtained two different molecular subtypes of sarcopenia: Cluster A (enriched in inflammatory pathways, e.g., IL6/TNF-α) and Cluster B (linked to mitochondrial dysfunction, e.g., PGC-1α suppression). These subtypes exhibited divergent clinical trajectories, suggesting subtype-specific management strategies—anti-inflammatory therapies for Cluster A and metabolic modulators for Cluster B. Validate subtype-associated prognosis and therapeutic responses in longitudinal cohorts. Decipher genetic/environmental drivers (e.g., epigenetic modifications, lifestyle factors) of subtype divergence. Develop subtype-targeted therapies (e.g., JAK/STAT inhibitors for Cluster A or mitophagy enhancers for Cluster B) to optimize outcomes.

Nonetheless, there were several limitations in the present work. First, we did not incorporate wet-lab experiments to validate the identified candidate biomarkers. Second, the limited sample size may introduce statistical biases. Third, the absence of long-term follow-up data impedes the evaluation of the predictive model’s stability and the reliance on single-source data restricts cross-platform applicability.

By integrating multi-omics data, this study systematically deciphered molecular features associated with sarcopenia and constructed a novel, effective diagnostic model. The results help understand the disease pathogenesis and highlight potential directions for developing precision therapeutic interventions. However, more experiments are needed for verifying the robustness of the conclusions, and expanding the cohort size is essential to improve generalizability. Future efforts should prioritize multidisciplinary collaborations to accelerate translational innovation and advance clinical applications in this field.

## Author contributions

YW, SG and YL analyzed and organized the data; YW and SG wrote and revised the manuscript; YZ designed, revised and supervised the study. All of the authors reviewed and approved the final manuscript.

## Data availability

The data that support the findings of this study are available upon request from the corresponding author.

